# A multi-parent advanced generation inter-cross population for genetic analysis of multiple traits in cowpea (*Vigna unguiculata* L. Walp.)

**DOI:** 10.1101/149476

**Authors:** Bao-Lam Huynh, Jeffrey D. Ehlers, Maria Munoz-Amatriain, Stefano Lonardi, Jansen R. P. Santos, Arsenio Ndeve, Benoit J. Batieno, Ousmane Boukar, Ndiaga Cisse, Issa Drabo, Christian Fatokun, Francis Kusi, Richard Y. Agyare, Yi-Ning Guo, Ira Herniter, Sassoum Lo, Steve I. Wanamaker, Timothy J. Close, Philip A. Roberts

## Abstract

Development and analysis of Multiparent Advanced Generation Inter-Cross (MAGIC) populations have been conducted with several crop plants to harness the potential for dissecting the genetic structure of traits and improving breeding populations. We developed a first MAGIC population for cowpea (*Vigna unguiculata* L. Walp.) from eight founder parents which are genetically diverse and carry many abiotic and biotic stress resistance, seed quality and agronomic traits relevant to cowpea improvement in sub-Saharan Africa (SSA) where cowpea is vitally important in the human diet and in local economies. The eight parents were inter-crossed using structured matings to ensure the population would have balanced representation from each of the founder parents, followed by single-seed descent, resulting in 365 F8 recombinant inbred lines (RILs) each carrying a mosaic of genome blocks contributed from all founders. This was confirmed by SNP genotyping with the cowpea Illumina 60K iSelect BeadArray. Following filtering to eliminate duplicates, sister lines and accidental selfing events, a core set of 305 F8 RILs was chosen as the primary population. The F8 lines were on average 99.74% homozygous while also diverse in agronomic traits including flowering time, growth habit, maturity, yield potential and seed characteristics across environments. Trait-associated SNPs were identified for most of the parental traits. Loci with major effects on photoperiod sensitivity and seed size were also verified by genetic mapping in biparental RIL populations. The distribution of recombination frequency varied considerably between chromosomes, with recombination hotspots distributed mostly in the telomeric regions. Due to its broad genetic base, this cowpea MAGIC population promises breakthroughs in genetic gain and high-resolution genetic mapping for gene discovery, enhancement of breeding populations and, for some lines, direct releases as new varieties.

## Introduction

Cowpea (*Vigna unguiculata* L. Walp) is a highly nutritious warm-season grain legume crop vitally important for food security in Africa where it provides a primary source of protein that complements cereals in the diet (Ehlers and Hall 1997; Kudre *et al.* 2013) and fodder for livestock. However, in the Sudano-Sahel region of West Africa typical smallholder farmer cowpea grain yields are only 10-20 % of known yield potential (Widders 2012). Biotic stresses caused by insect pests, and diseases caused by pathogens, the parasitic weed *Striga gesnerioides* and nematodes, and abiotic stresses from heat, drought and low-fertility soils are primary constraints to cowpea grain production. Many of these problems also affect cowpea production in parts of southern Europe, Asia, Australia, Latin America, and southern United States (Ehlers and Hall 1997; Huynh *et al.* 2013a). Development of cowpea cultivars that tolerate or resist these constraints will increase yield and reduce costly chemical based crop-protection inputs and promote human and environmental health, thus directly benefitting resource-poor farmers.

The greatest opportunity to increase cowpea grain yields lies in the rich genetic variation within this diploid (2n = 22) species as it has numerous resistance and tolerance traits to combat biotic and abiotic stresses (Huynh *et al.* 2013b; Muchero *et al.* 2013). Many of these traits have been genetically mapped using QTL discovery and mapping approaches (OuÉdraogo *et al.* 2002b; Muchero *et al.* 2010; Muchero *et al.* 2011; Lucas *et al.* 2012; OuÉdraogo *et al.* 2012; Pottorff *et al.* 2012; Pottorff *et al.* 2014; Huynh *et al.* 2015; Huynh *et al.* 2016). Cowpea has the capacity to produce grain under magnitudes of water stress that would render comparable crops unproductive (Ewansiha and Singh 2006), yet significant differences in drought tolerance exist among cowpea lines at different stages of growth (Watanabe *et al.* 1997; Mai-Kodomi *et al.* 1999a). For example, there are significant phenotypic differences in ability to survive vegetative-stage drought stress (Mai-Kodomi *et al.* 1999b; Muchero *et al.* 2013), providing opportunity for cowpea breeders to incorporate early-season drought tolerance into improved varieties. Among genotypes exhibiting seedling drought tolerance, two types of responses were observed by Mai-Kodomi et al. (1999a). Type 1 response plants ceased all growth and conserved moisture in all plant tissues, thereby allowing subsequent recovery of the entire shoot upon re-hydration. In contrast, type 2 response involved plants mobilizing moisture from lower leaves to sustain growth of new trifoliates, with rapid senescence of unifoliates at the onset of water-stress conditions. Mid- and late-season drought stresses have received considerable attention, given their negative effects on yield parameters (Hall *et al.* 2003; Padi 2004; Dadson *et al.* 2005). On a physiological level, osmotic adjustment, carbon isotope discrimination, transpiration, assimilation rates, and stomatal conductance in cowpea have been studied in detail (Hussain *et al.* 1999; Anyia and Herzog 2004; Odoemena 2004). In many cases, however, results were inconclusive or no meaningful differentiation between genotypes was achieved. Morphological investigations have tended to focus on root-related parameters where genotypes were compared for rooting depth and relative root biomass (Matsui and Singh 2003; Ogbonnaya *et al.* 2003). Phenologically, flowering and maturation times have been investigated for drought escape-related strategies (Gwathmey and Hall 1992). Early maturing varieties may be able to complete their reproductive cycle in time to escape late-season drought (Grantz and Hall 1982; Ehlers and Hall 1997), but these varieties were sensitive to mid-season drought (Thiaw *et al.* 1993). Early flowering coupled with the delayed leaf senescence trait, which later promotes survival during mid-and late-season drought, allowing plants to produce a second flush of pods, offers the greatest potential for managing both mid-and late-season drought conditions (Gwathmey and Hall 1992). Association mapping identified multiple loci with pleiotropic effects on drought-related traits in cowpea across experiments in West Africa under limited water conditions (Muchero *et al.* 2013). Because drought tolerance is a complex trait, its genetic improvement combined with selection for biotic resistance would need a systematic breeding strategy involving multiple trait donors.

Development of multi-parent advanced generation intercross (MAGIC) populations, termed by Cavanagh *et al.* (2008), provides a state-of-the-art approach to advancing plant population resources for genetic analysis and breeding. It involves inter-mating multiple elite parents for several cycles followed by single-seed descent, resulting in recombinant inbred lines (RILs) each carrying a mosaic of genome blocks contributed from all founders. Development and analysis of MAGIC populations have been undertaken in a few crops including wheat, rice and chickpea due to the potential of this approach for dissecting genetic and genomic structure (Huang *et al.* 2015). The goal of the current work was to develop an 8-parent MAGIC population for cowpea using founder parents that are highly diverse and carry many key traits relevant to cowpea production in SSA. In this paper, we report on the development, genetic analysis and validation of this new genetic resource using high-density marker genotyping platforms (Muchero *et al.* 2009a). Due to its broad genetic base, the cowpea MAGIC population provides opportunities for increasing genetic gain and QTL/gene discovery in cowpea and related species.

## Materials and Methods

### Choice of parents

The eight cowpea parents used in the original crosses were elite cultivars and breeding lines selected based on their high genetic diversity characterized by genotyping with 1,536 genome-wide gene-based SNP markers (Muchero *et al.* 2009a; MuÑoz-AmatriaÍn *et al.* 2017). In addition, they were chosen because collectively they carry multiple biotic and abiotic stress resistance and tolerance traits relevant to SSA (Table 1). SuVita 2, also known as ‘Gorom’, a local landrace in Burkina Faso, is resistant to the parasitic weed Striga (OuÉdraogo *et al.* 2002b) and the fungal pathogen *Macrophomina phaseolina* (Muchero *et al.* 2011). CB27, a California blackeye cultivar bred by University of California–Riverside (UCR) (Ehlers *et al.* 2000), is heat tolerant (Lucas *et al.* 2013a) and highly resistant to root-knot nematodes (Huynh *et al.* 2016), Fusarium wilt disease (Pottorff *et al.* 2012; Pottorff *et al.* 2014), and foliar thrips (Lucas *et al.* 2012). IT93K-503-1, a breeding line from the International Institute of Tropical Agriculture (IITA) breeding nursery in Nigeria, is drought tolerant (Muchero *et al.* 2009b), and resistant to root-knot nematodes (Huynh *et al.* 2016), *M. phaseolina* (Muchero *et al.* 2011), and Fusarium wilt (Pottorff *et al.* 2014). The other five parents (IT89KD-288, IT84S-2049, IT82E-18, IT00K-1263, and IT84S-2246) are also breeding lines from IITA, Nigeria; they also carry combinations of key traits including grain quality and resistance to root-knot nematode, Striga, Fusarium, viruses, and bacterial blight (Table 1).

**Table 1.** MAGIC founder parents and their traits relevant to SSA and other production areas

| Name | Source<sup>a</sup> | Agronomic trait | Resistance or tolerance trait |
| --- | --- | --- | --- |
| SuVita 2 | INERA | High yielding under drought in Senegal, Burkina Faso and Mozambique; large dark-brown seed | Drought tolerant, resistant to Striga, foliar thrips and *Macrophomina* disease |
| CB27 | UCR | High yielding under drought in Mozambique; large black-eye seed; photoperiod insensitive; erect growth habit; early maturing | Heat tolerant, resistant to root-knot nematode, Fusarium wilt, and foliar thrips |
| IT93K-503-1 | IITA | High yielding under drought in Senegal; brown-eye seed; stay-green under drought | Drought tolerant, resistant to nematodes, Fusarium wilt, and *Macrophomina* |
| IT89KD-288 | IITA | High yielding under drought in Burkina Faso and Nigeria; brown-eye seed; photoperiod sensitive | Root-knot nematode resistant |
| IT84S-2049 | IITA | High yielding under drought in Burkina Faso; brown-eye seed; erect growth habit | Resistant to aphid, bacterial blight, viruses, root-knot nematode |
| IT82E-18 | IITA | High yielding under drought in Mozambique; early maturing, light-brown seed; photoperiod insensitive | Broadly adapted, resistant to root-knot nematode |
| IT00K-1263 | IITA | High yielding under drought in Mozambique and Nigeria; dark-brown seed; stay-green under drought | Resistant to Striga, aphid, fusarium wilt, root-knot nematode |
| IT84S-2246 | IITA | High yielding under drought in Burkina Faso and Mozambique; dark-brown seed | Resistant to aphid, bacterial blight, viruses, root-knot nematode |

<sup>a</sup> INERA: Institut de l'Environnement et des Recherches Agricole, Burkina Faso; UCR: University of California – Riverside, United States; IITA: International Institute of Tropical Agriculture, Nigeria.

### Population development

The eight parents were inter-mated using a mating strategy described in Cavanagh et al. (2008) with some modifications. In spring 2010, initial crosses were made between 4 pairs of founder parents (IT89KD-288 x IT84S-2049, CB27 x IT82E-18, SuVita 2 x IT00K-1263, and IT84S-2246 x IT93K-503-1, designated as A x B, C x D, E x F, and G x H, respectively) to produce 2-way F1s. In spring 2011, reciprocal 4-way crosses were made between 2 pairs of the 2-way F1s to produce 4-way F1s. In fall 2011, 330 pair crosses were made between 330 4-way F1 plants of the pedigrees ABCD and 330 4-way F1 plants of the pedigrees EFGH to produce 331 8-way F1s. Single seed descent (SSD) was then applied for each unique 8-way F1 until the F8 generation. For each F8 RIL, seeds from the F8 single plant were harvested and maintained as an original seed stock (F8:9). The F8:9 seeds were then increased in bulk to make F8:10 seeds for phenotyping.

### SNP genotyping

The F1 progeny from 2-way crosses were verified by genotyping their F2 seeds (up to 21 seeds per cross) with 89 parent-unique SNPs using the kompetitive allele-specific polymerase chain reaction (KASP) cowpea assay (LGC Genomics Ltd., Hoddesdon, UK) (Semagn *et al.* 2014), which was converted from the 1536 SNP Illumina GoldenGate Assay developed by Muchero et al. (2009a). True F1 plants were confirmed when polymorphic markers were found segregating in the corresponding F2 progeny.

The F8 single plants derived from 8-way crosses were genotyped with 51,128 SNPs using the Illumina iSelect BeadArray (MuÑoz-AmatriaÍn *et al.* 2017). A core set of MAGIC RILs was selected through the following consecutive steps: (1) Lines carrying non-parental alleles or excess numbers of heterozygous and ambiguous genotypes were excluded; (2) Based on parent-unique SNPs, lines that did not carry male-parent alleles (i.e., selfing) at the 4-way or 8-way crosses were also excluded; (3) Among true 8-way RILs and eight parents, genetic similarities were measured using the allele-sharing method (BOWCOCK *et al.* 1994) with the software GGT 2.0 (van Berloo 2008), from which phylogenetic relationships were generated using the neighbor-joining method (Saitou and Nei, 1987) and visualized using the software MEGA 5.05 (Tamura et al., 2011); and (4) For each set of genetically identical RILs (similarity 0.99 or higher), the line with the lowest number of ambiguous genotypes was retained in the core set for further analyses.

### MAGIC phenotyping

The MAGIC RILs and parents were screened for photoperiod sensitivity under long-daylength conditions during summer, from June (14.5 hours) to September (12.8 hours), at the UCR Citrus Experiment Station, California (UCR-CES, 33.97°N, 117.34°W). In 2015, each MAGIC RIL and parent was planted in one row of 0.76 m wide and 5.5 m long at a density of 12 seeds per meter using a tractor-mounted planter. The field was watered to capacity before and after planting up to 100 days using furrow irrigation. The experiment was repeated in 2016 but under restricted irrigation. The field was watered to capacity before planting, and then irrigation was withheld until the end of trial. For each line in both trials, calendar days to flowering were determined when 50% of plants in the plot flowered.

The population was also screened under short daylength conditions during autumn, from September (12.8 hours) to December (9.9 hours), at the Coachella Valley Agricultural Research Station, California (CVARS, 33.52°N, 116.15°W). In 2015, the population was planted in two blocks receiving different watering regimes (full irrigation and restricted irrigation) and separated by a 6-row buffer (5 m). In each block, each MAGIC RIL and parent was planted in one row of 0.76 m wide and 3.5 m long at a density of 12 seeds per meter using a tractor-mounted planter. The field was watered to capacity before and after planting using subsurface drip irrigation. After two weeks when the seedlings were well established, the irrigation was withheld in the restricted-irrigation block until maturity, whereas in the full-irrigation block the rows were watered to capacity up to 100 days after planting. In 2016, the two experiments (full irrigation and restricted irrigation) were repeated on adjacent field blocks at CVARS. For each line in four experiments, calendar days to flowering were determined when 50% of plants in the plot flowered. Plant growth habit was measured 40 days after planting using a visual rating scale from 1 to 6 based on the angles formed between primary branches and the main stem: (1) Acute erect, branches form angles less than 45° with the main stem, (2) Erect, branching angles between 45° – 90° with the main stem, (3) Semi-erect, branches perpendicular to the main stem but not touching the ground, (4) Intermediate, lower branches touching the ground, (5) Semi-prostrate, lower branches flat on the ground but the main stem standing upright, and (6) Prostrate, the entire plant flat and spreading on the ground. Days to maturity were determined when 95% of pods in the plot had dried. At maturity, the plants in each plot were cut at the lower stems and machine-threshed for measurement of plot yield and 100-seed weight.

For each set of repeated trials at UCR-CES and CVARS, analysis of variance (ANOVA) was performed with the software GenStat version 11 (PAYNE *et al.* 2008). Factors in the ANOVA model were lines and block, with each field site receiving one watering treatment considered as a block. Broad-sense heritability was estimated based on the variance component attributable to variation among lines (VG) and residual variation (VE) (h^2^ = VG/(VG + VE)). Nonparametric Spearman’s rank correlation analysis was also used to examine the consistency in genotypic ranking of the same lines between environments.

### Association mapping

Polymorphic SNPs (success rate > 90% and minor allele frequency > 0.05) with known positions across 11 cowpea pseudomolecules (Lonardi *et al.* 2017) were used for genetic mapping, with a new chromosome numbering convention based on synteny between cowpea and common bean. Genome-wide association studies (GWAS) were performed with the software TASSEL 5.0 (Bradbury *et al.* 2007) using the mixed linear model (MLM) function that incorporated population principal components and kinship analyses. A LOD threshold of 4 was used to indicate QTL significance.

To verify QTL detected for photoperiod sensitivity in the MAGIC population, a biparental mapping population including 92 F8-derived F9 RILs from a cross between the non-photoperiod sensitive parent CB27 and the photoperiod sensitive IT97K-556-6 was screened under long-daylength conditions at UC Riverside in 2016. Each RIL and parent were planted in one row 0.76 m wide and 5.5 m long at a density of 12 seeds per meter using a tractor-mounted planter. The planting time and experimental conditions were similar to the MAGIC phenotyping trial in 2015. For each plot, days to flowering were determined when 50% of plants in the plot flowered. The biparental RIL population was genotyped with the 51,128 SNP Illumina iSelect BeadArray that was used to genotype the MAGIC population. Construction of genetic maps and QTL analysis were performed with the software QTL IciMapping 4.0 (Meng *et al.* 2015) using the Inclusive Composite Interval Mapping (ICIM) method (Wang 2009).

### Recombination analysis

Based on physical positions of polymorphic SNPs on 11 cowpea pseudomolecules, pair-wise linkage distances (in centimorgans) between every two adjacent SNPs were estimated as *d* = [(*recombinants*/*n*) x 100], where *recombinants* are MAGIC lines that carry SNP genotypes that are not present in any of the eight parents (Supplemental File S1), and *n* is the number of MAGIC lines carrying valid SNP genotypes. Using a sliding window of 2 Mb with 1 Mb increments along each chromosome, recombination rates (cM/Mb) were calculated as the linkage distance divided by the physical distance between the first and the last SNP of each window. The recombination-rate variation was visualized by plotting the estimated recombination rate for every 1-Mb increment along the 11 chromosomes.

### Resource and data availability

The MAGIC core set and their 8 founder parents are available on request at the IITA, Ibadan, Nigeria and University of California Riverside, USA cowpea germplasm banks. QTL information for photoperiod sensitivity and seed size is provided in Supplemental Files S2 and S3, respectively. Genotypic and phenotypic data used in GWAS and biparental mapping are included in Supplemental Files S4 and S5, respectively.

## Results

### MAGIC development and genotyping

In total 365 MAGIC F8 RILs were generated from 330 unique 8-way crosses. Among these, 29 crosses produced two or more F8 RILs, which were sister lines separated from earlier generations. These lines were purposely created to maintain the population size. Reciprocal crosses were made at the 4-way and 8-way cycles, resulting in RILs with different maternal parents, including CB27 (225 lines), IT89KD-288 (111 lines), Suvita 2 (9 lines), and IT84S-2246 (5 lines). There were 15 lines with illegible pedigrees on tags that were bleached by sunlight and moisture in the greenhouse.

Genotyping with the 51,128-SNP Illumina iSelect BeadArray resulted in 36,346 SNPs that were polymorphic between the 8 parents (68.26%). Among these, 11,848 SNPs were parent-unique, each of which could distinguish one parent from the other 7 parents. Based on parent-unique SNPs, 15 RILs lacked male-parent alleles at the 2-way/4-way intercrosses and probably resulted from accidental selfing during artificial pollination. Three RILs were found to carry non-parental alleles in that they were heterozygous at SNP loci that were monomorphic between the 8 parents. Except for five RILs with more than 10% of heterozygosity, the rest of the population had a low level of heterozygosity, ranging from 0 to 3.33%. There were 8 RILs each showing very similar SNP genotypes (more than 99%) to another RIL, and these were considered as redundant duplicates. After excluding lines with duplicates, selfing errors, non-parental alleles and excess heterozygosity, a core set of 305 MAGIC RILs derived from 305 unique 8-way crosses was selected for further analysis. These RILs appeared highly diverse and clustered uniformly relative to their eight parents, among which IT89KD-288, IT84S-2246 and IT97K-503-1 were closer to each other than the other parent to parent relationships which were dispersed throughout the population structure (Fig. 1).

**Figure 1.**
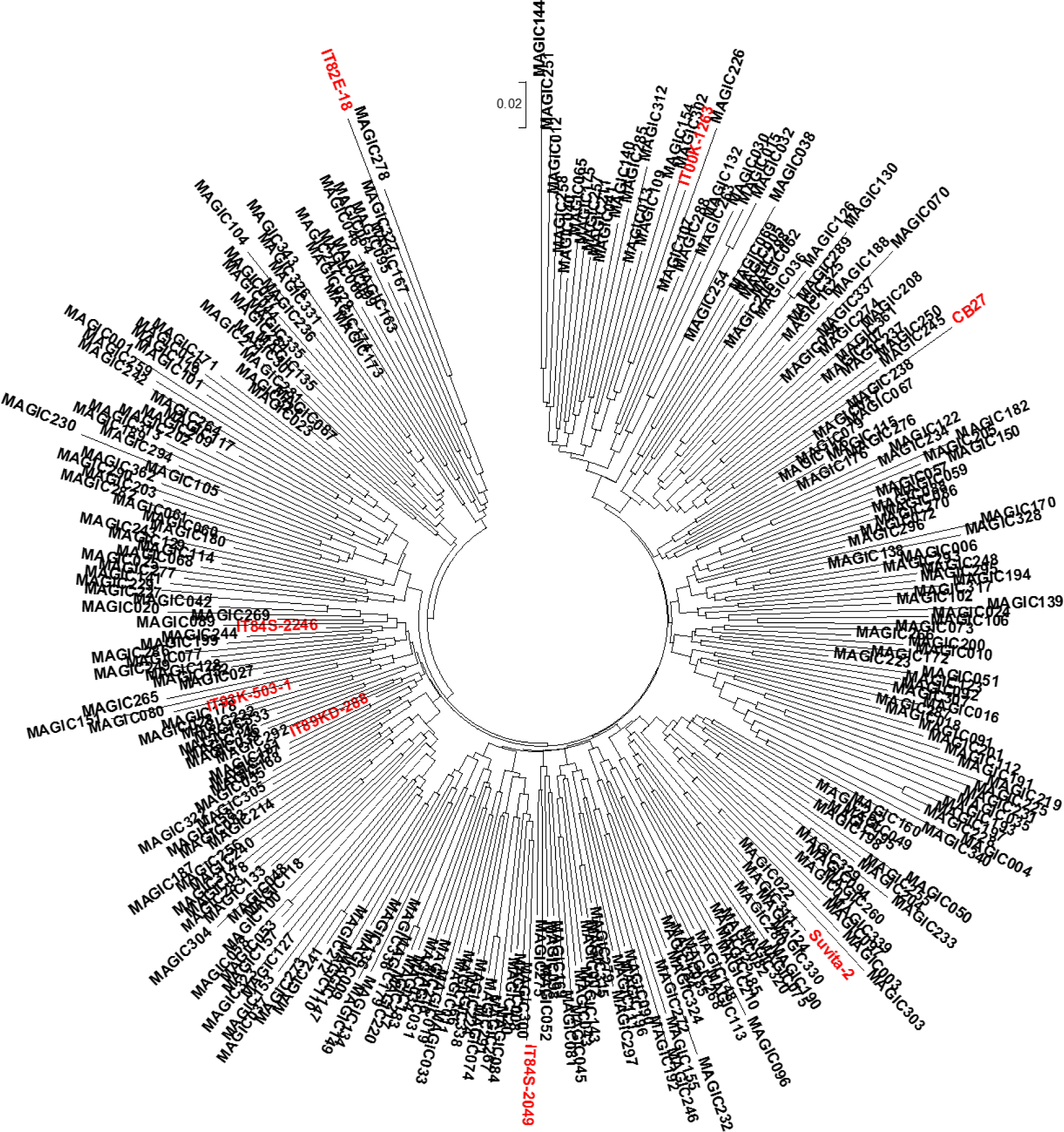
Phylogenetic relationships among the 305 F8 RILs of the cowpea MAGIC core set and eight parents (in red) based on 11, 848 parent-unique SNPs

### Phenotypic variation in the MAGIC population

The F8 lines were highly diverse in morphological traits including flowering time, growth habit, flower color, leaf shape, and seed characteristics (size, shape, color and texture) (Fig. 2). The flowering time varied widely in the population under both long- and short-daylength conditions (Fig. 3). The genotypic differences in flowering time were quite stable across contrasting watering regimes in each daylength condition, with broad-sense heritability estimated as 0.77 and 0.71 at UCR-CES (long daylength) and CVARS (short daylength), respectively. There was a significant correlation (*r* = 0.63, *P* < 0.001) in phenotypic ranking between the long- and short-daylength conditions, although the absolute flowering time varied considerably among lines. At UCR-CES (long daylength), the population started flowering as early as 43 days after planting, but there were many lines with delayed flowering beyond 60 days after planting (Fig. 3a and 4a). In contrast, under short daylength at CVARS, the population started flowering as early as 34 days after planting and the population completed flowering within another month (63 days) (Fig. 3b and 4b). Among the parents, CB27 (36 and 44 days) was the earliest to flower while IT89KD-288 (46 and 88 days) was the most delayed in both environments (CVARS and UCR-CES, respectively).

**Figure 2.**
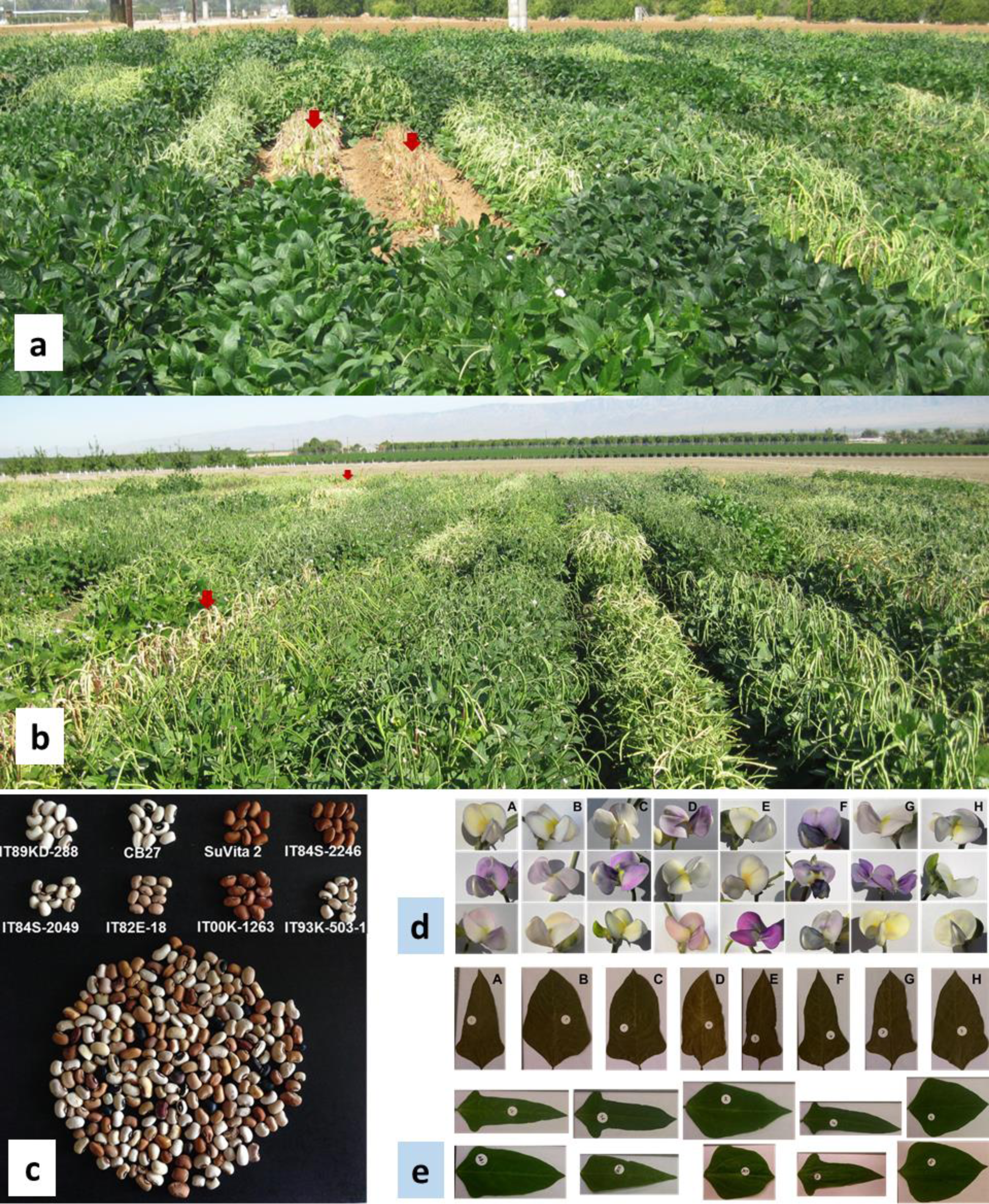
Morphological variation in the cowpea MAGIC population: Plant appearance at 65 days after planting under (a) long-daylength conditions at UCR-CES in 2015 and (b) short-daylength conditions at CVARS in 2016, both under full irrigation; (c) seed appearance, (d) flower color and (e) leaf shape of parents (top panel) and a representation of MAGIC F8 RILs (in lower part of 2c, each seed is from a different F8 RIL). In 2a and 2b, red arrows indicate examples of lines that matured earlier than other lines. In 2d and 2e, parent codes are: A, IT89KD-288; B, IT84S-2049; C, CB27; D, IT82E-18; E, Suvita-2; F, IT00K-1263; G, IT84S-2246; H, IT93K-503-1.

**Figure 3.**
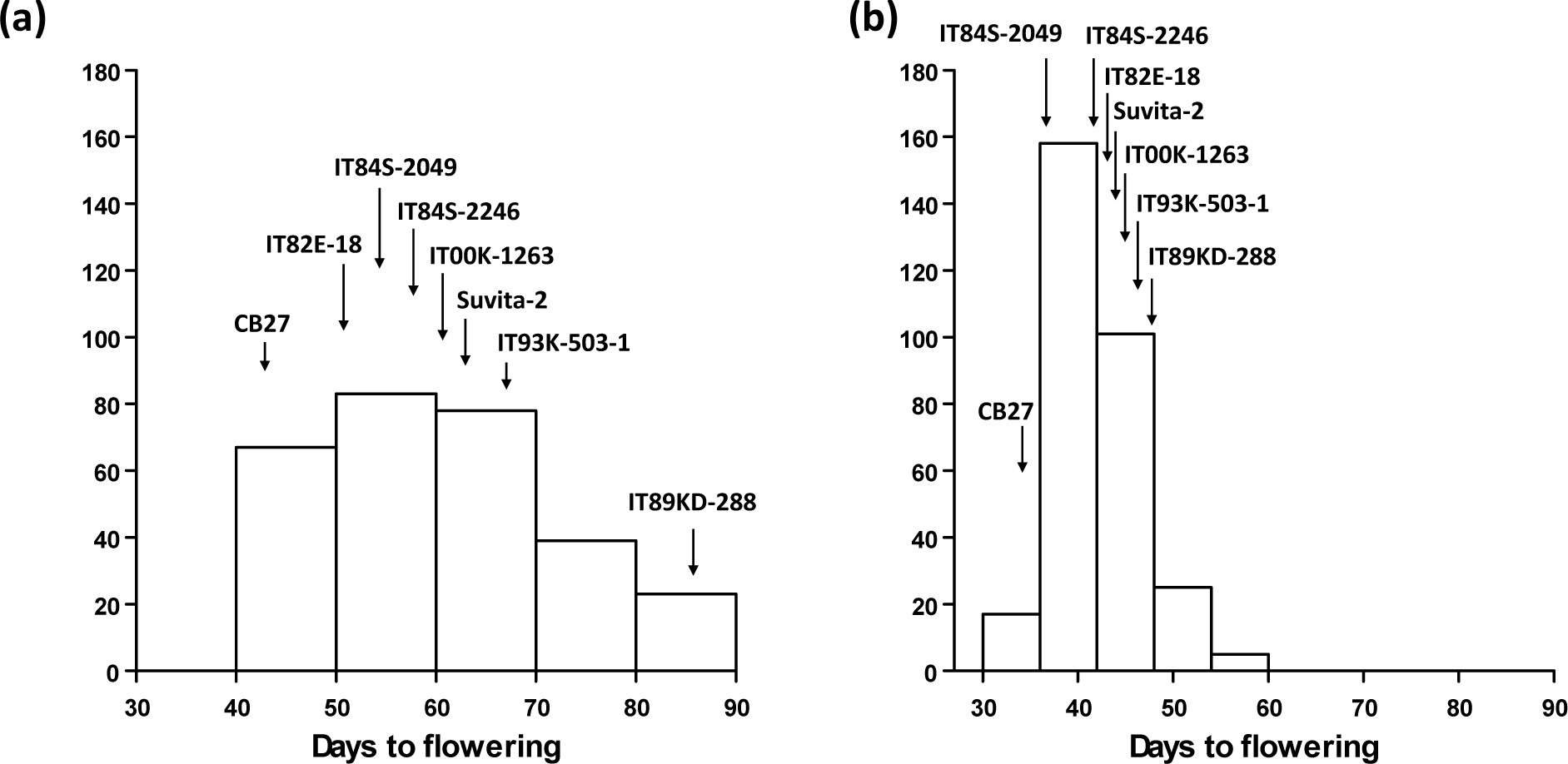
Variation in flowering time measured in the MAGIC core set and eight parents grown under (a) long-daylength conditions at UCR-CES and (b) short-daylength conditions at CVARS. Mean flowering time values for each line were derived from two experiments at UCR-CES and four experiments at CVARS during 2015-2016.

None of the MAGIC RILs or parents showed prostrate growth habit. Under full irrigation, the majority of MAGIC RILs had a growth habit ranging from semi-erect to erect under both short- and long-daylength conditions (Fig. 4). There was significant but moderate correlation in the growth habit scores between the two daylength conditions (*r* = 0.55, *P* < 0.001), with about 55% of the lines showing consistent growth habit between the two environments. Lines with semi-prostrate growth habit under long-daylength became intermediate or semi-erect type when grown under short-daylength condition. Among the parents, CB27 (acute erect) and IT84-2049 (erect), IT89KD-288 (semi-erect) and Suvita-2 (semi-erect) maintained their growth habit in both short- and long-daylength conditions under the full-irrigation regime. Under restricted irrigation, the MAGIC RILs and parents mostly showed erect or acute erect growth.

**Figure 4.**
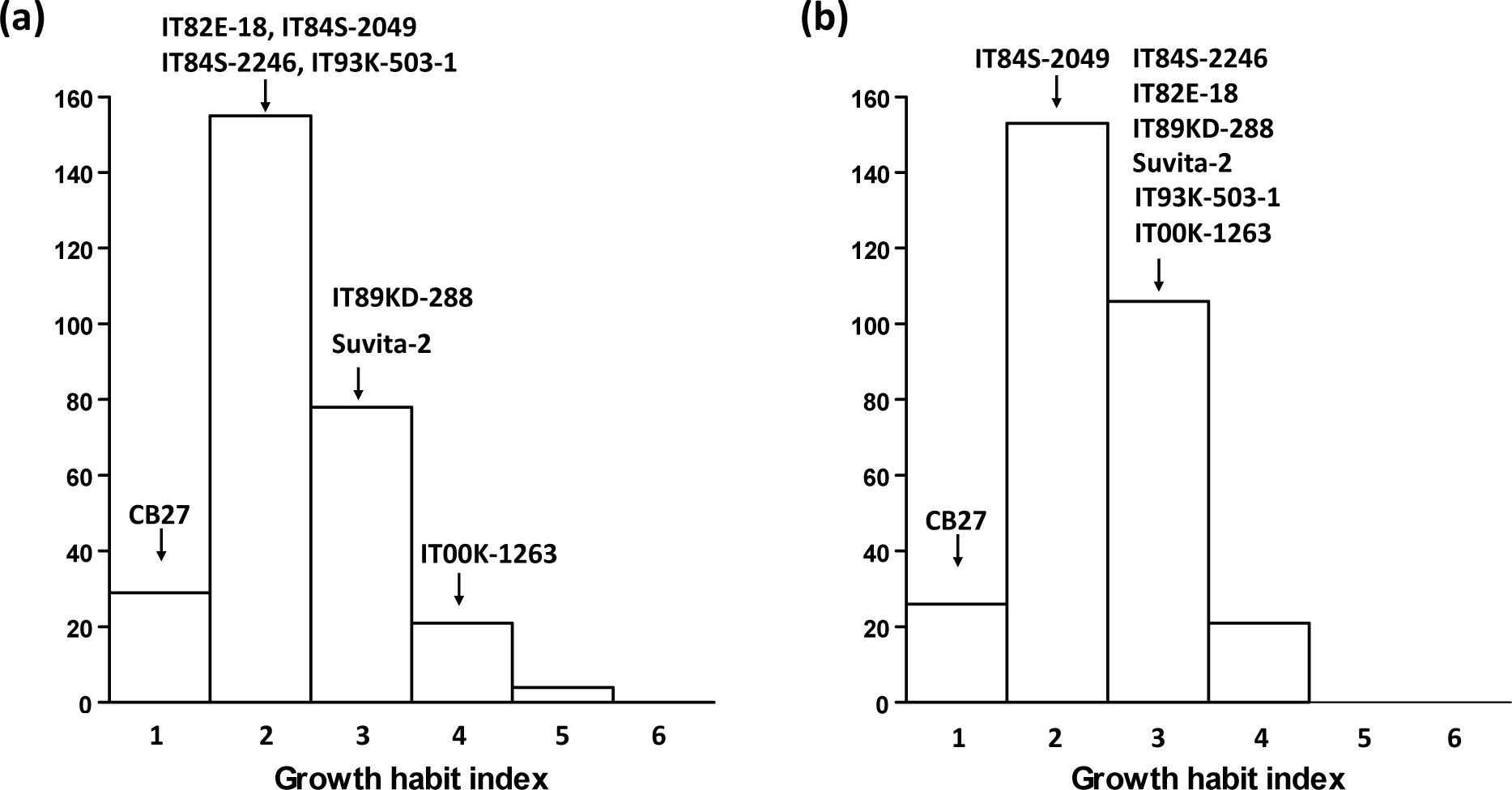
Variation in plant growth habit measured in the MAGIC core set and eight parents grown under full irrigation with different photoperiod conditions: (a) long-daylength at UCR-CES in 2015 and (b) short-daylength at CVARS in 2015 and 2016 (means of 2 years). Growth habit index:(1) acute erect, (2) erect, (3) semi-erect, (4) intermediate, (5) semi-prostrate, and (6) prostrate.

Maturity varied considerably in the MAGIC population grown under different watering regimes at CVARS in 2015 and 2016. Broad-sense heritability was estimated as 0.47, with significant but moderate correlations (*r* = 0.53, *P* < 0.001) existing in the phenotypic ranking between the two watering conditions (normal and restricted irrigation). Transgressive segregation was also observed. Some lines were fully mature as early as 60 days after planting under both watering regimes, while others were still green and kept producing pods after 120 days under restricted irrigation in 2015, including two parents (IT00K-1263 and IT93K-503-1) and 66 MAGIC RILs (21% of the population).

Grain yield and seed size also varied considerably under both water-restricted and full irrigation conditions at CVARS (Fig. 5). The plants generally produced much higher yield and developed larger seeds under full irrigation compared to water-stress conditions. Seed size appeared much more stable in the genotypic ranking than grain yield, with broad-sense heritability estimated as 0.76 and 0.30, respectively. Transgressive segregation was observed for both traits. Approximately 11% of MAGIC RILs yielded higher than all parents under restricted irrigation conditions. Among the parents, CB27 consistently had the highest yield and seed size across the two environments.

**Figure 5.**
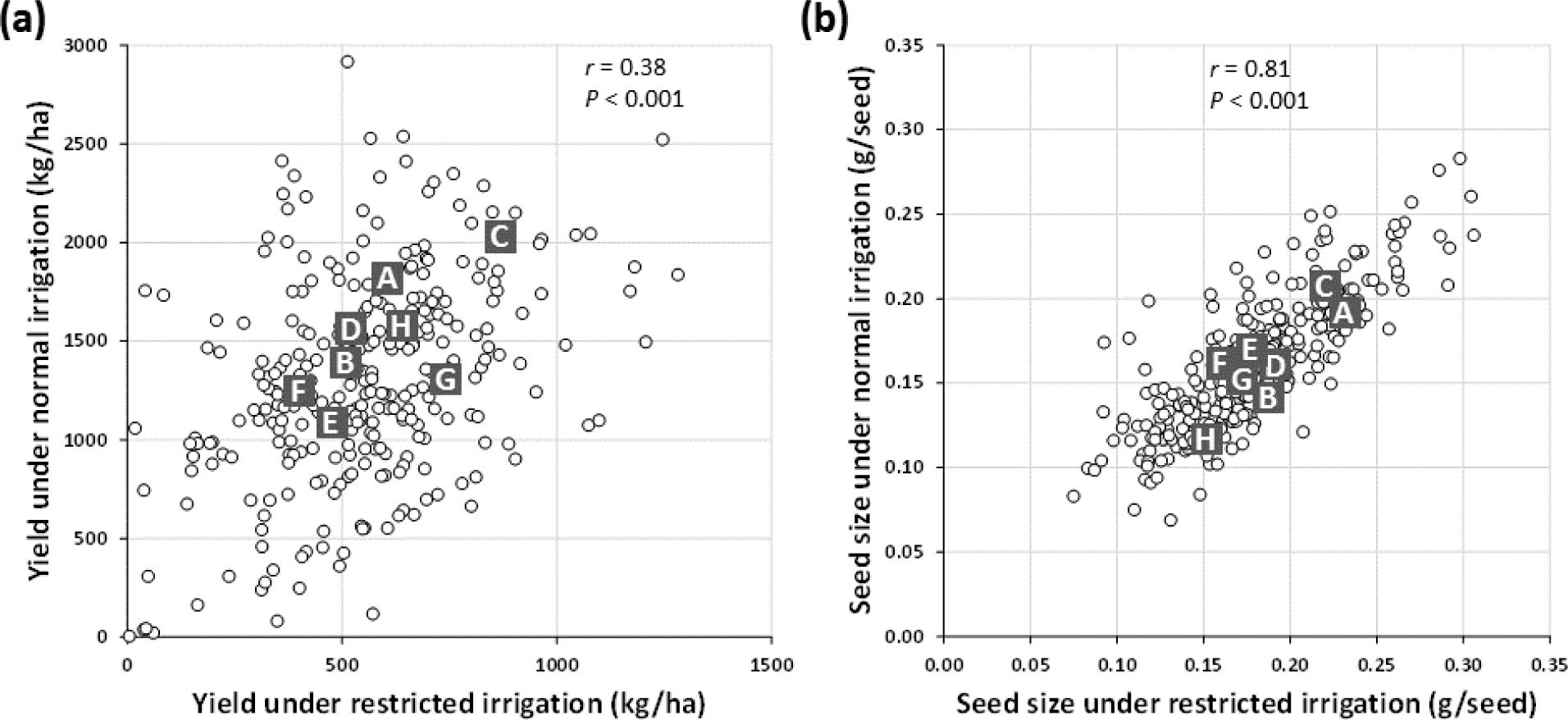
Variation in (a) dry grain yield and (b) seed size measured in the 305-RIL MAGIC core set and parents grown under restricted and normal irrigation regimes at CVARS in 2015 and 2016. Each dot represents a MAGIC RIL based on mean phenotypic values of two years of data. Labeled solid squares are MAGIC parents (A: IT89KD-288, B: IT84S-2049, C: CB27, D: IT82E-18, E: Suvita-2, F: IT00K-1263, G: IT84S-2246, H: IT93K-503-1).

### Detection of marker-trait associations

From the 36,346 polymorphic SNPs, after removing those with MAF ≤ 0.05 and successful calling rate ≤ 90%, the remaining 33,768 SNPs with known positions on 11 cowpea pseudomolecules were used in GWAS. The MLM analysis consistently detected three clusters of SNPs on chromosomes 5, 9 and 11 significantly associated with photoperiod sensitivity expressed in the MAGIC population grown under long-daylength conditions at UCR-CES in 2015 (full irrigation, Fig. 6a) and 2016 (restricted irrigation, Fig. 6b). Markers with major effects were located on chromosome 9 (max LOD = 9.3), explaining up to 15% of total phenotypic variance, with favorable (earlier flowering time) alleles contributed from CB27 and IT82E-18. Markers with smaller effects were located on chromosomes 5 and 11 (max LOD = 5.8), explaining up to 9% of total phenotypic variance. Under the short-daylength condition, the significant peaks on chromosomes 5 and 9 remained, albeit with smaller effects (max LOD = 6.5, explaining 9% of phenotypic variance) while the peak on chromosome 11 was not significant (Fig. 6c and 6d).

**Figure 6.**
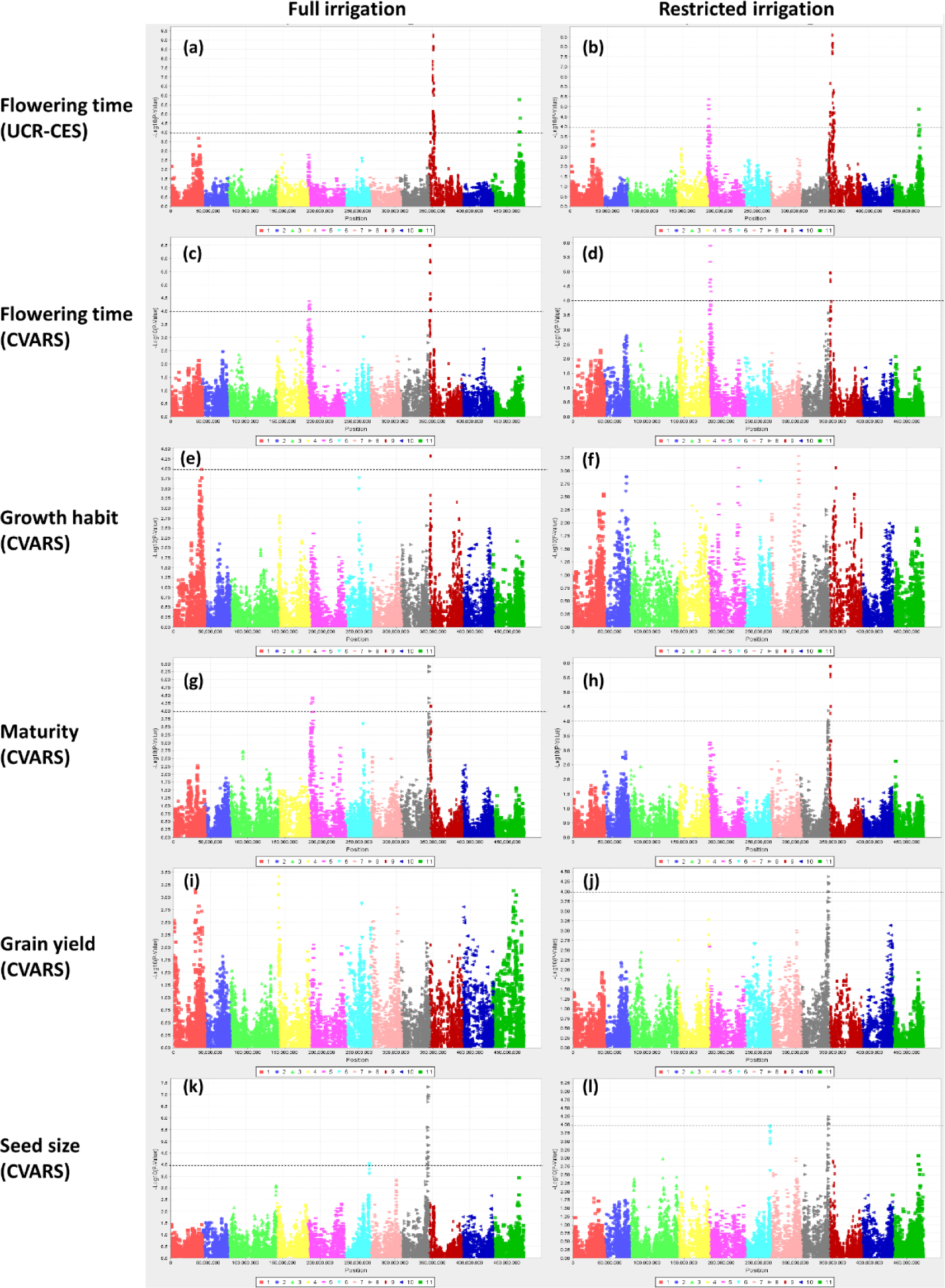
GWAS Manhattan plots depicting chromosomal regions associated with variation in agronomic traits measured in the MAGIC population grown under full irrigation (left) and restricted irrigation (right) with long-daylength conditions at UCR-CES (a-b) and short-daylength conditions at CVARS (c-l). Phenotypic values at CVARS were means of two years of data, except for growth habit which was not measured under restricted irrigation in 2016. The dashed lines correspond to the significance threshold −log_10_(0.0001).

Markers significantly associated with plant growth habit were identified on distal regions of chromosomes 1 and 9 based on data from short-daylength and full irrigation experiments at CVARS (Fig. 6e). Markers of highest significance (max LOD = 4.3) explained up to 9% of total phenotypic variance, with favorable alleles (more erect growth) contributed from IT89-KD288 and CB27. This region also coincided with markers affecting flowering time under the short-daylength condition (Fig. 6d), with the more erect-growth alleles associated with earlier flowering time. No significant association was found for growth habit under restricted irrigation.

Marker peaks on chromosomes 5, 8 and 9 with significant or marginal effects on maturity also coincided with marker peaks affecting flowering time under short-daylength conditions at CVARS (Fig. 6g and 6h). For grain yield under restricted irrigation at CVARS, a significant peak was detected, located on chromosome 8 (max LOD = 4.4, explaining 6% of phenotypic variance) (Fig. 6j) while no significant markers were identified for grain yield under normal irrigation (Fig. 6i). In contrast, markers with major effects were identified for seed size on chromosome 8 based on data from both water–restricted and full-irrigation conditions (Fig. 6k and 6l). Markers of highest significance (max LOD = 7.3) explained up to 11% of total phenotypic variance, with favorable (larger seed) alleles contributed by IT82E-18 and IT00K-1263. These markers also flanked the region previously mapped for seed size using a biparental RIL population (CB27 x IT82E-18), with the favorable allele contributed from IT82E- 18 (LUCAS *et al.* 2013b) (Supplemental File S2). The other region with minor effects on seed size under both water regimes was found located on chromosome 6 (max LOD = 4.0), explaining 5% of total phenotypic variance, with favorable alleles contributed from IT89KD-288, Suvita-2, IT84S-2246 and IT93K-503-1.

In general, markers significantly associated with agronomic trait determinants appeared to distribute around distal regions of chromosomes. Neither maternal effect nor its interaction with trait-associated SNPs for each agronomic trait were significant.

### Validation of photoperiod QTL in biparental RILs

Flowering time varied widely in the CB27 x IT97K-556-6 RIL population grown under long-daylength condition at UCR-CES in 2016 (Fig. 7). CB27 began flowering 44 days after planting while IT97K-556-6 delayed flowering until after 70 days. A major QTL for flowering time was detected on linkage group 9 (LOD = 7.8, explaining 30% of phenotypic variance) (Fig. 8). The favorable (early flowering) allele was contributed from CB27. SNP markers flanking this QTL were 2_04691 and 2_00735 (and their co-segregating SNPs), which were found to be located in the same region as the major QTL detected in the MAGIC population grown under the same long-daylength condition in 2015 (Supplemental File S3).

**Figure 7.**
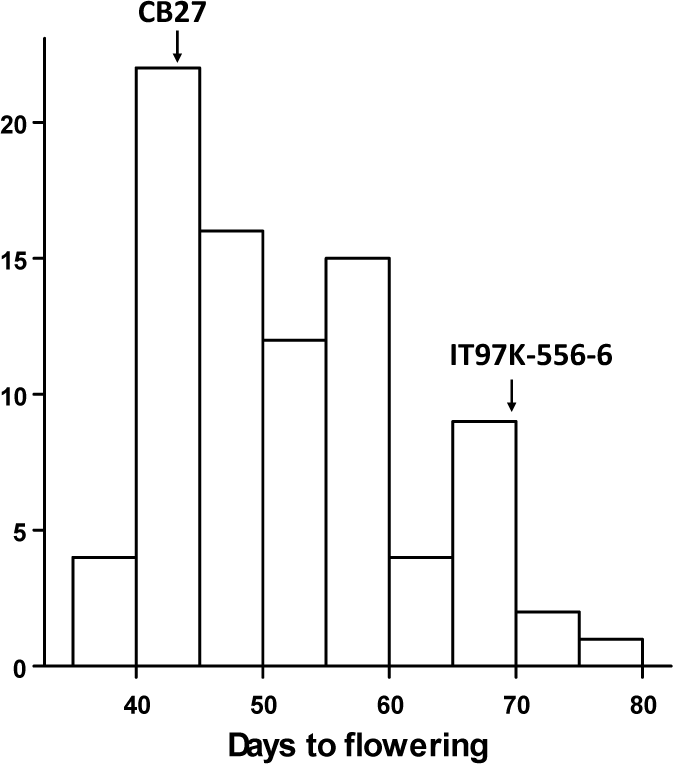
Variation in flowering time expressed in the CB27 x IT97K-556-6 biparental RIL population grown under long-daylength conditions at UCR-CES in 2016.

**Figure 8.**
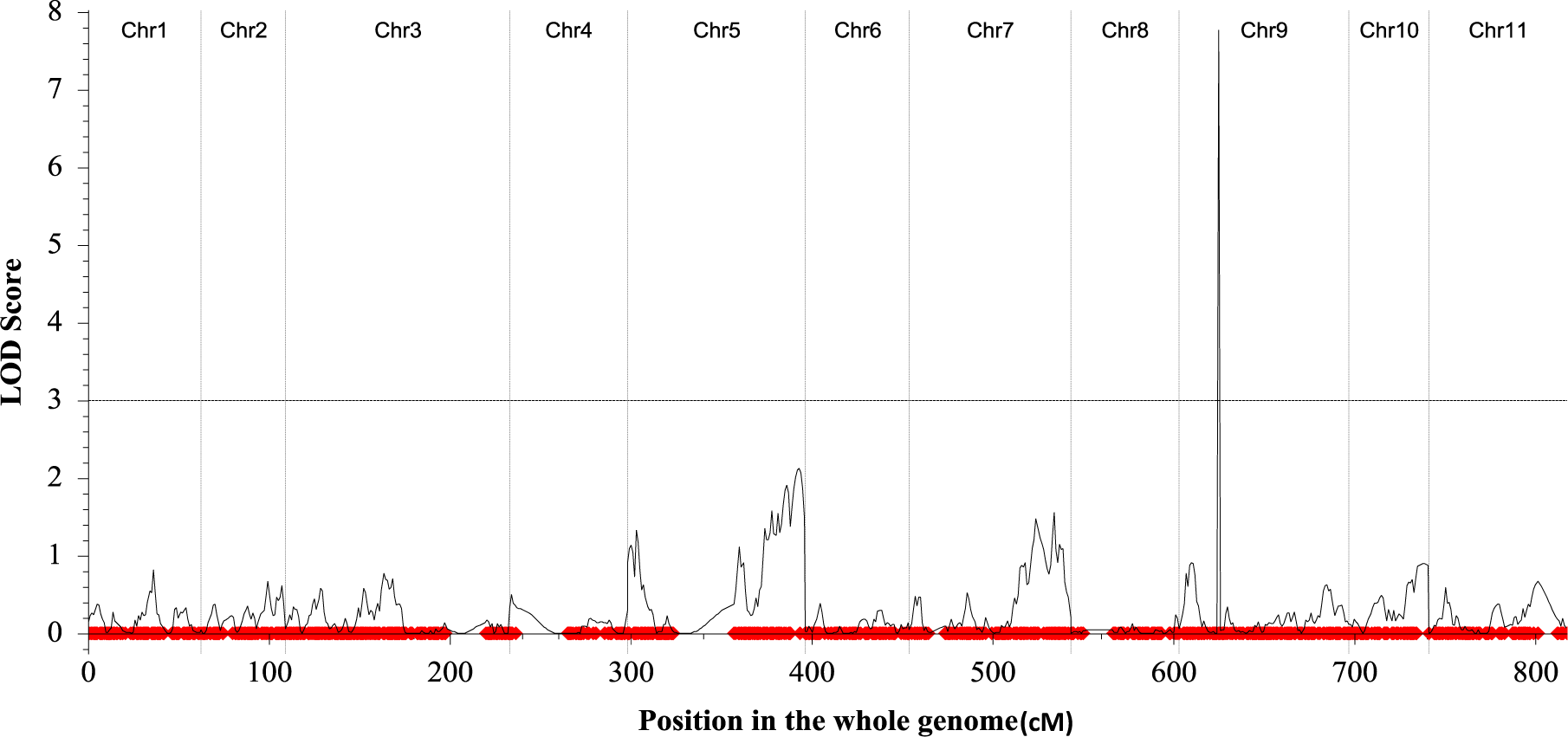
Chromosomal regions associated with variation in flowering time measured in the CB27 x IT97K-556-6 biparental RIL population under long-daylength conditions at UCR-CES in 2016. The LOD peak on linkage group 8 is flanked by SNP markers 2_10023 and 2_04691, which are in the same region of the major QTL detected in the MAGIC population (see Fig. 6a).

### Recombination rate variation

Based on recombination analysis of 33,768 polymorphic SNPs, the crossovers appeared to distribute throughout the MAGIC genome, at an average of 1.43 cM/Mb, and more frequently on or near the telomeric distal regions of chromosomes (Fig. 9), where most trait-associated SNPs were detected by GWAS (Fig. 6). Several recombination hot-spots with more than 5 cM/Mb were detected on the distal long arms of chromosomes 1, 2, 5 and 11, while fairly large disequilibrium blocks also were found on most chromosomes. On average, chromosome 3 had the highest recombination rate (1.76 cM/Mb) while chromosome 10 had the lowest (0.88 cM/Mb).

**Figure 9.**
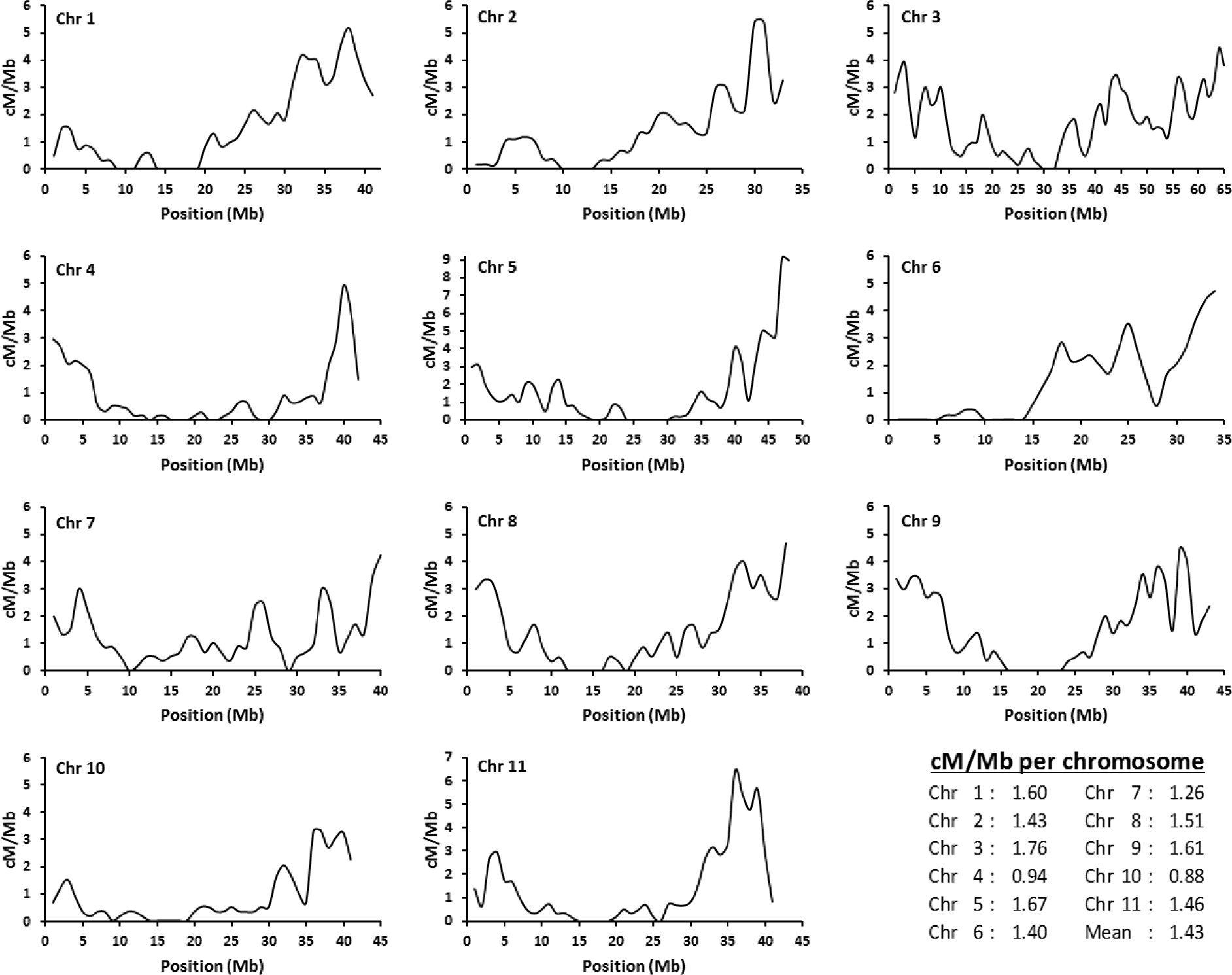
Recombination rate (cM/Mb) variation along 11 cowpea chromosomes measured in the 8-way cowpea MAGIC population using a sliding window of 2 Mb with 1 Mb increments.

## Discussion

### MAGIC development

The wide phenotypic variation with significant transgressive segregation observed in the cowpea MAGIC population indicates that genome regions from parents were highly recombined in the RILs. In fact, the population was developed in a way that maximized genetic variation. At the 2- way crosses, plants in each F1 set were heterozygous and homogeneous because the 8 founder parents were highly inbred lines. However, the F1s derived from the 4-way crosses segregated and exhibited significant variation. To capture as much variation as possible, we performed more than 300 pair-wise crosses, also including reciprocal pairs, between different 4-way F1 individuals. In addition, there was no intended selection for any trait during the SSD process. The plants were grown in UCR greenhouses with optimal temperature, fertilizer, irrigation and pest and disease management. In some cases, the plants were grown during long-daylength conditions in summer, but the photoperiod sensitive lines which failed to become reproductive in the summer were maintained and allowed to set flowers and pods later in the autumn when the daylength shortened, to avoid selection against photoperiod sensitivity. There was also no selection for preferable seed characteristics, plant type or yield components. This blind SSD process therefore helped create the high diversity in morphological and agronomic traits that we observed in this MAGIC population (Fig. 2-5).

The genetic integrity of the cowpea MAGIC population was also confirmed by the results of high-density SNP genotyping. We used 89 parent-unique SNP markers from the Illumina GoldenGate Assay (Muchero *et al.* 2009a) to validate true 2-way F1 crosses to avoid possible mistakes from the early stage of MAGIC development. We then used 11,848 parent-unique SNPs from the recently developed Illumina iSelect 60K SNP assay (MuÑoz-AmatriaÍn *et al.* 2017) to confirm true 8-way RILs and to eliminate those that appeared to be selfed at the 4-way or 8-way crosses. The SNP genotyping also identified lines with non-parental alleles, identical SNP genotypes, or excess heterozygosity. Fortunately, very few of these unexpected lines were found (16 out of 365 lines), some of which were replaced by sister lines that were purposely developed as backups. By removing all erroneous lines and keeping one RIL from each unique 8-way cross, we created a MAGIC core set of 305 RILs that are highly homozygous and genetically distinct from each other and from the eight parents (Fig. 1). As such, they can serve as permanent genetic materials for use in replicated phenotyping trials. Despite its compact size relative to MAGIC populations developed for *indica* rice (2000 lines) (Bandillo *et al.* 2013) and winter wheat (1091 lines) (Mackay *et al.* 2014), the cowpea 305 RIL MAGIC core set is less management-intensive and easier to phenotype, especially under field conditions, while still capable of accurately detecting trait-associated markers as discussed in the following section.

### MAGIC genetic analysis

Based on SNP genotyping, the 36,346 markers segregating in the cowpea MAGIC population were almost double those identified in every bi-parental RIL population genotyped with the same SNP array (MuÑoz-AmatriaÍn *et al.* 2017). This is attributable to the 8 founder parents having been chosen on the basis of phenotypic and genetic diversity found in earlier studies. The parents were high yielding under drought in one or more countries, resistant to different biotic stress factors (Table 1), and represented the two major gene pools of cowpea centered in West Africa and southeastern Africa (Huynh *et al.* 2013a). By applying multiple 2-way, 4-way and 8-way intercrosses from those founders, plus 8 generations of single seed descent for over 300 independent 8-way pair crosses, one would expect more recombination events to occur in the MAGIC than in bi-parental RILs. However, it is difficult to measure accurately the number of crossovers between two SNP markers due to a lack of parent-specific alleles at each locus. At each SNP marker, one allele represents one or more parents, and the alternative allele represents the other parents, so in some cases it is impossible to identify the actual parent carrying the allele at that locus if multiple parents bear a common allele. The recombination rate estimated in this study was based only on the obvious recombinants between two SNPs (Supplemental File S1) and thus may underestimate their true genetic distance. Despite this, the current estimated recombination rate was still relatively high and varied considerably along 11 cowpea chromosomes (Fig. 9), supporting a high level of crossing over as expected in the MAGIC population. The pattern of variation along chromosomes appears consistent with information reported for *Drosophila* (Hey and Kliman 2002) and human genomes (Mcvean *et al.* 2004) where the recombination was often elevated near telomeres and suppressed near centromeres.

The relatively high recombination rates near the telomeres imply that haplotype variation occurs at a high level, thereby facilitating random association needed for QTL detection. This is in accordance with our GWAS analysis showing that trait-associated SNPs were mostly distributed near the telomeric regions (Fig. 6). In addition, markers with major effects detected in the cowpea MAGIC population for photoperiod sensitivity and seed size were verified by bi-parental genetic mapping, indicating that the MAGIC core set is effective for mapping genome regions harboring major QTLs. This MAGIC core set comprised individuals which were carefully selected based on genome-wide SNP diversity excluding sister lines and duplicates, so interference by kinship and population structure would not be a problem during association analysis. The population size of 305 highly homozygous lines also provides sufficient internal replications of marker alleles needed for marker-trait association; at a parent-unique locus, approximately one-eighth of the population (~38 individuals or 12.5%) carry the parent-unique allele. The challenge for QTL detection in the MAGIC population, however, is a lack of donor-specific markers flanking the QTL in question, whose alleles cannot distinguish the QTL donor from other parents. In the case of the major photoperiod QTL, it was fortunate that favorable alleles in the region were specific to the non-photoperiod donors CB27 and IT82E-18 (Supplemental File S3). For other trait-associated SNPs with lower LOD scores, their true effect might be diluted by ‘hidden’ unfavorable alleles during GWAS analysis. Enriching such regions with more parent-unique markers would help improve the efficiency of QTL mapping.

### Genetic improvement perspectives

The strong transgressive segregation observed for agronomic traits provides opportunities for selecting MAGIC lines that outperform the parents. Selecting for large seed size, which is preferred by consumers in SSA, would be straightforward because the trait appeared highly heritable (Fig. 5b). In contrast, selecting for higher yield will be more difficult given its relatively low heritability (Fig. 5a); based on the pattern of variation in yield under restricted versus full irrigation, it may be more effective to select for high yield under drought stress in which at least 11% of the RILs yielded better than the 8 MAGIC parents. These lines probably carry a combination of different drought-tolerance genes contributed from multiple parents, because the parents are known to yield well under drought conditions in different African countries (Table 1). MAGIC lines that are not photoperiod sensitive could be grown widely across seasons and regions with different latitudes. Lines that flower early may escape damage by flower/pod feeding insects and abiotic stress such as heat and terminal drought in certain environments. Lines with exceptionally early or delayed crop senescence are suitable for production systems requiring single or double flushes of pods, respectively. Plants with acute erect growth could support heavy pod load, allow more leaf area to capture sunlight for photosynthesis, and support high plant population densities to increase yield under monocropping. Since the eight parents also vary in resistance to many major insects and diseases (Table 1), the MAGIC population will segregate for many biotic stress resistance traits and also contain lines with unique and novel combinations of defense genes. Therefore, phenotypic screening of the MAGIC population for those traits will enable genetic mapping and identification of lines carrying favorable trait combinations for selecting cultivars in target environments.

For the longer term, the cowpea MAGIC population can also benefit breeding programs by providing valuable pre-breeding resources. QTLs detected in the MAGIC population combined with existing knowledge of QTL regions and haplotypes can be applied to develop novel combinations of QTLs through intercrossing best MAGIC RILs, providing super trait-donor lines for use in breeding programs. QTLs for many key traits were already mapped in bi-parent and diversity populations where certain MAGIC parents were used in the crosses, such as seed size (Lucas *et al.* 2013b), heat tolerance (Lucas *et al.* 2013a), drought tolerance (Muchero *et al.* 2013), resistance to foliar thrips (Lucas *et al.* 2012), aphids (Huynh *et al.* 2015), Fusarium wilt disease (Pottorff *et al.* 2014), root-knot nematodes (Huynh *et al.* 2016), ashy stem blight or charcoal rot disease caused by *Macrophomina phaseolina* (Muchero *et al.* 2011), viruses (OuÉdraogo *et al.* 2002a), the parasitic weed *Striga gesnerioides* (OuÉdraogo *et al.* 2012), and root architecture (Burridge *et al.* 2017). It is therefore possible to track positive haplotypes contributed by different MAGIC parents in each MAGIC line and then intercross best lines to develop target ideotypes. The strategy would be similar to the multi-parent advanced generation recurrent selection (MAGReS) approach proposed recently by Huang et al. (2015), except that (1) prior knowledge of QTL information from cowpea bi-parental mapping will be utilized, (2) the MAGIC RILs selected for intercrosses are more advanced (F8), and (3) the selection can be targeted using both QTL haplotypes and predicted breeding values based on genome-background diversity. The resulting MAGReS lines will be fixed for positive haplotypes at known QTLs and carry additional recombinations in other unknown loci conferring high grain yield. They can thus be a valuable resource both for genetic improvement (as super trait donors or new cultivars) and for detecting novel QTLs when combined with the current MAGIC RIL set.

## Supplementary Materials

Supplemental File S1: Illustration of recombinant identification

Supplemental File S2: Major QTL region associated with seed size

Supplemental File S3: Major QTL region associated with photoperiod sensitivity

Supplemental File S4: Genotypic and phenotypic data used in GWAS

Supplemental File S5: Genotypic and phenotypic data used in biparental mapping

## Acknowledgements

This work was supported in large part by grants from the Generation Challenge Programme of the Consultative Group on International Agricultural Research to PAR, JDE and TJC, with additional support from the USAID Feed the Future Innovation Lab for Collaborative Research on Grain Legumes (Cooperative Agreement EDH-A-00-07-00005) to PAR and TJC, the USAID Feed the Future Innovation Lab for Climate Resilient Cowpea (Cooperative Agreement AID-OAA-A-13- 00070) to TJC and PAR, and the NSF-BREAD (Advancing the Cowpea Genome for Food Security) to TJC, PAR, SL, MMA and BLH. The authors thank Tra Duong, Hyun Park Kang, Eric Castillo, Jasmine Gracin-Dixon, Uriah Dixon, Mitchell Lucas and Savanah St. Clair for technical assistance. The authors also thank Peggy Mauk, Vince Samons, and staff at UC Riverside Agricultural Operations and Coachella Valley Agricultural Research Station for managing field trials.

